# Hitchhiking in space: ancestry in adapting, spatially-extended populations

**DOI:** 10.1101/084426

**Authors:** Brent Allman, Daniel B. Weissman

## Abstract

Selective sweeps reduce neutral genetic diversity. In sexual populations, this “hitchhiking” effect is thought to be limited to the local genomic region of the sweeping allele. While this is true in panmictic populations, we find that in spatially-extended populations the combined effects of many unlinked sweeps can affect patterns of ancestry (and therefore neutral genetic diversity) across the whole genome. Even low rates of sweeps can be enough to skew the spatial locations of ancestors such that neutral mutations that occur in an individual living outside a small region in the center of the range have virtually no chance of fixing in the population. The fact that nearly all ancestry rapidly traces back to a small spatial region also means that relatedness between individuals falls off very slowly as a function of the spatial distance between them.

## Introduction

In large populations even a fairly low rate of selective sweeps is sufficient to reduce diversity across most of the genome via hitchhiking (Gillespie, 2000; Weissman and Barton, 2012). Most modeling of the effects of hitchhiking on diversity has considered well-mixed populations. However, the effects are potentially quite different in spatially-extended populations with only short-range dispersal, because instead of quickly fixing through logistic growth, sweeps must spread out in a spatial wave of advance over the whole range (Fisher, 1937). Barton et al. (2013) recently showed that this increase in the time to sweep tends to reduce the size of the genomic region over which diversity is depressed by a sweep. While the effect of sweeps on genetic diversity at linked loci is therefore reduced by spatial structure, we show here that collective effect of sweeps on the diversity at *unlinked* loci can be much stronger than in panmictic populations. Surprisingly, this effect is dependent on the geometry of the range - it only appears for realistic range shapes with relatively well-defined central regions, not for the perfectly symmetric idealizations of ring-shaped and toroidal ranges often used in theoretical models. In particular, we find that probability of fixation of an allele can be strongly position-dependent, with alleles near the center of the range orders of magnitude more likely to fix than those at typical locations. This is because all individuals trace most of their ancestry, even in the not-too-distant past, to individuals living in the center, which also causes far-away individuals to be much more closely related to each other than they would be in the absence of the unlinked sweeps, with relatedness falling off only as a power law of distance rather than exponentially.

## Model

We wish to find the expected number of copies that an allele found in an individual at spatial position *x* will leave far in the future, i.e., its reproductive value (Barton and Etheridge, 2011), which we denote *ϕ*(x). Equivalently, *ϕ*(x)*ρ*(x), where *ρ* is the population density, is the probability density of a present-day individual’s ancestor being at location *x* at some time in the distant past. Maruyama (1970) showed that in the absence of selection, *ϕ*(x)≡1 regardless of the details of the population structure, as long as dispersal does not change expected allele frequencies. Here we show that this result does not extend to populations undergoing selection. Populations living in perfectly symmetric ranges (circles in one dimension, tori in two) necessarily have *ϕ*(x)≡1, but when this symmetry is broken, recurrent sweeps can make reproductive value strongly dependent on spatial position, with high *ϕ* in a small region in the center of the range and very small *ϕ* everywhere else.

We consider a population with uniform, constant density *ρ* distributed over a *d*-dimensional range with radius *L*, with uniform local dispersal with diffusion constant *D*, i.e., 2*D* is the mean squared displacement after unit time. We assume that selective sweeps with advantage s occur in the population at a rate Λ per generation, originating at points uniformly distributed over time and space, and at loci uniformly distributed over the genome. As long as the density is sufficiently high (*ρ* ≫ (*s*/*D*)^*d*/2^ /*s*, Nagylaki (1978); Barton et al. (2013)), they will spread roughly deterministically in waves with speed 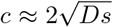 with characteristic wavefront width 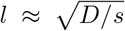 (Fisher, 1937), which we take to be much smaller than the range size, *l* ≪ *L*. (However, even for fairly large densities, the stochastic corrections to *c* can be substantial; see Eq. 16 in the Methods.) We assume that Λ is low enough compared to the frequency of outcrossing, *f*, and the average number of crossovers per outcrossing, *K*, that the waves do not interfere with each other. For well-mixed populations, this means that Λ ≪ *f K* (Weissman and Barton, 2012); we are currently preparing a manuscript in which we show that spatially-extended populations have nearly the same limit on Λ, up to logarithmic factors. The definitions of symbols are collected in Table 1.

**Table 1:** Symbol definitions

| Symbol | Definition |
| --- | --- |
| $d$ | Number of spatial dimensions (1 or 2) |
| $\rho$ | Density of individuals (individuals / (distance)<sup>$d$</sup>)* |
| $L$ | Radius of range* |
| $D$ | Dispersal constant* |
| $f$ | Frequency of outcrossing |
| $K$ | Expected number of crossovers per outcrossing |
| $s$ | Selective advantage of adaptive alleles |
| $\Lambda$ | Frequency of selective sweeps |
| $c \approx 2\sqrt{Ds}$ | Expected rate of advance of a sweeping beneficial allele |
| $l \approx \sqrt{D/s}$ | Characteristic width of the wave of advance of a sweep |
| $\phi(x)$ | Reproductive value of individuals at location $x$ |
| $\psi(x)$ | Identity-by-descent between individuals separated by $x$ |
| *In continuous space. In the corresponding model of discrete demes on a square lattice, $\rho$ is the deme size, $L$ is the radius in demes, and $D$ is half the rate of migration into a deme, i.e., $d$ times the migration rate between a pair of neighboring demes. | |

## One and two dimensions

We consider both one-dimensional ranges (lines with length 2*L*) and two-dimensional ranges. In two dimensions, the shape of the range will have some effect on many of our results; however, as long as the shape is fairly “nice”, with a clear center and single characteristic length scale *L*, this effect will be modest. We will therefore ignore it for simplicity. For our purposes, the main difference between one and two dimensions will be in the density of individuals a distance *x* from the center, *ρ*(*x*). Since we are assuming a uniform spatial density, in one dimension this is just *ρ*, a constant. In two dimensions, however, we must account for the fact that there is more area at larger *x*, and thus *ρ*(*x*) ≈ 2*πxρ*. (Necessarily, *ρ*(*x* > *L*) = 0 in both one and two dimensions.)

## Results

In spatially-extended populations, genetic hitchhiking not only changes the frequency of neutral alleles, but also shifts their distribution in space. To see this, consider the ancestry of a lineage going backward in time, so that sweeps appear as receding waves. When one passes over the focal lineage, it “pulls” it back towards origin of the sweep at the same speed *c* as the wave. If there is no recombination, the lineage will necessarily be pulled all the way back to the origin (i.e., all present-day individuals necessarily descend from the original mutant at the swept locus), but if recombination is frequent, the lineage will be pulled only a short distance before recombining out of the wave and stopping. If recombination occurs at rate *r*, then we expect that the lineage will remain in the wavefront for a time of 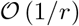 before recombining, and therefore be pulled a distance of ~ *c/r* towards the origin of the sweep. For most positions in most realistic range shapes, sweeps tend to arrive (forward in time) from the direction of the center of the range, and so pull the ancestry back towards the center; the collective effect of many sweeps is then to concentrate the ancestry in the center.

To make this description more quantitative, it will be convenient to classify sweeps based on their genetic map distance *r* to the focal locus. We will refer to sweeps at *r* ≪ *s* as *tightly-linked* and those at *r* ≫ *s* as *loosely-linked.* Barton et al. (2013) found that a tightly-linked sweep pulls a lineage a distance that is approximately exponentially distributed with mean *c/r*, going backward in time, with an upper cutoff at the distance to the origin of the sweep. In this paper, we calculate the effect of a loosely-linked sweep and find that the lineage is only pulled an expected distance *c/2r* (see Methods). To calculate the net effect of hitchhiking on a locus over time, we need to integrate over all sweeps occurring across the genome at different recombination fractions *r*. The 1/*r* dependence for the expected pull suggests that this net effect should be dominated by some combination of a few very tightly-linked sweeps and the many very loosely-linked sweeps (rather than the moderately-linked sweeps with *r* ~ *s*). This actually overstates the importance of tightly-linked sweeps, since the 1/*r* dependence has an upper cutoff for *r* ≾ *L/c*, and understates the importance of loosely-linked sweeps, since even if a sweep occurs very far away on the genome the recombination fraction cannot exceed *f*/2. Thus we expect that if the genome is sufficiently long (in a sense that will be made more precise below), the total average displacement of a typical locus will be dominated by loosely-linked sweeps. We will begin by focusing on this case, making the further approximation that most sweeps are not just loosely-linked but unlinked (*r* = *f*/2), as will be the case for even moderately long genomes, *K* ≿ 1. This case is also relevant for loci that are far from all loci undergoing selection, i.e., the ones whose evolution might be expected to depend only on demography. It also describes bacterial populations in which recombination primarily involves relatively short lengths of DNA, so that most pairs of loci in the genome recombine at roughly the same rate, as long as this recombination is still rapid relative to selection (the “quasisexual” case, Rosen et al. (2015)).

### The pull of unlinked sweeps

For a lineage a distance *x* from the center, there is an excess of approximately ~ Λ*x/L* sweeps per generation pulling it back toward the center, each of which pulls it an expected distance *c/f*. (Note that the effect of the upper cutoff on the displacement from these sweeps is negligible as long as *L ≫ c/f*.) The expected distance from the center therefore decays exponentially (backward in time) like

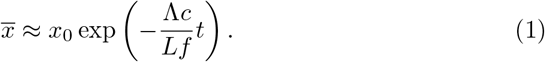

This implies that there is a characteristic concentration time *t*_con_ beyond which ancestry is significantly altered by the collective effect of unlinked sweeps:

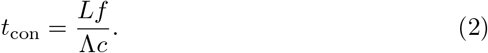

This deterministic move back to the center is opposed by dispersal, and also by the effect of occasional tightly-linked sweeps which pull the lineage a distance ~ *L*, effectively randomizing its position. The balances between these forces means that the ancestry of the population is not completely concentrated at the center of the range, but is instead distributed around it in some region of size ~ *x_c_*. Figure 1 shows this rapid concentration followed by a balance with dispersal and tightly-linked sweeps.

**Figure 1:**
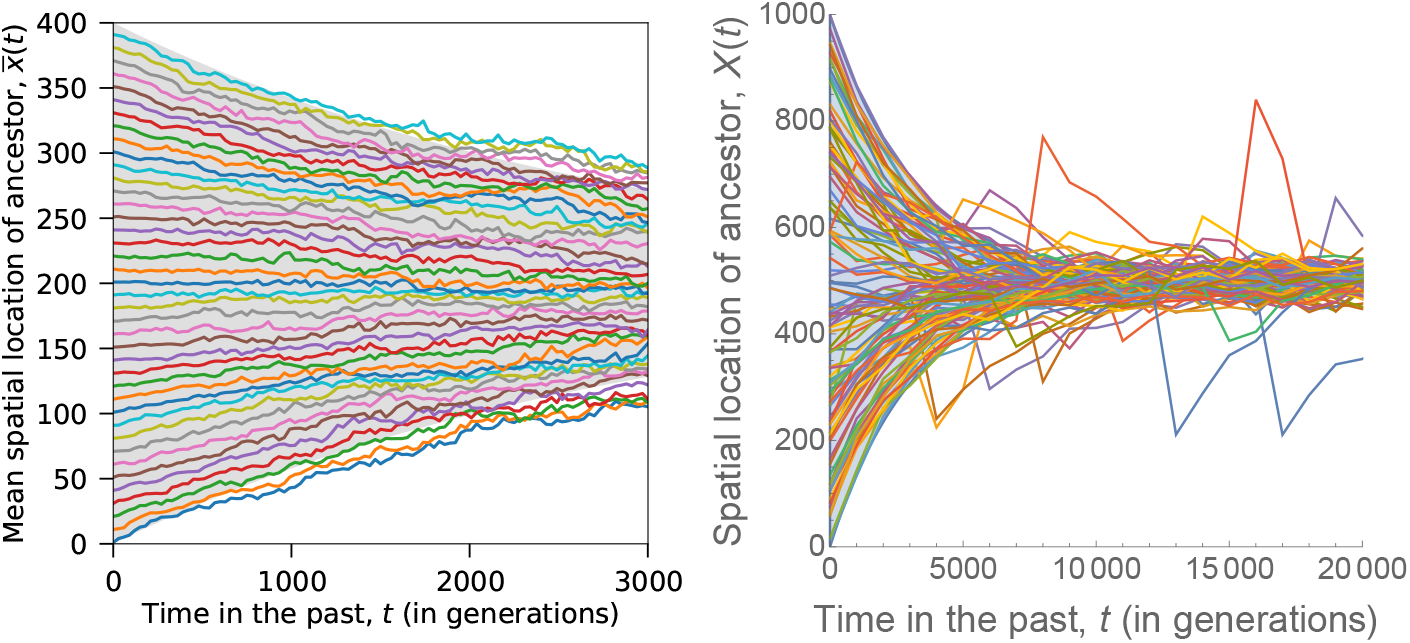
Tracing the ancestry of a neutral locus in a single individual back through time. Individuals throughout the range rapidly trace their ancestry back to a small region in the center of the range. Curves show simulations, while shaded regions show analytical predictions, Eq. 1. Left: Exact forward-time simulations. Each curve is the mean location of the ancestors of all individuals in a given present-day deme, averaged over three independent simulation runs. Parameters are *L* = 200, *s* = 0.05, *D* = 0.25, *f* = 1, *ρ* = 300, and Λ ≈ 0.6 (so that *t*_con_ ≈ 3000), with all loci unlinked. The discrepancy between the analytical prediction and the simulations at older times is an artifact caused by loss of resolution in the simulations as genetic diversity is exhausted (see Methods). Right: Approximate backward-time simulations. Each curve is an independent simulation of the ancestry of a single present-day individual. While the width of the central region is determined by a balance between dispersal and the pull of unlinked sweeps, the occasional excursions out of the center are due to hitchhiking on tightly-linked sweeps. Parameters are *L* = 500, *s* = 0.05, *D* = 0. 125, Λ = 1, *f* = 1, *K* = 300.

### Balance with dispersal

If tightly-linked sweeps are relatively rare, either because the overall rate of sweeps is low or because the focal locus lies in a region of the genome that is not undergoing much adaptation, the main balance will be between the diffusive effect of dispersal and the pull of unlinked sweeps. In this case, the position of the ancestry is an Ornstein-Uhlenbeck process, i.e., if we denote the position of the ancestral lineage *t* generations in the past by *X_t_*, it evolves backward in time as:

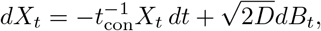

where *B_t_* is a Brownian motion. By Fick’s first law of diffusion, the diffusive flux of ancestry is –*D*∇*ϕ*(*x*). In the stationary state this must exactly cancel the deterministic pull of unlinked sweeps, so far in the past the distribution of ancestry is normal and concentrated in the center of the range according to:

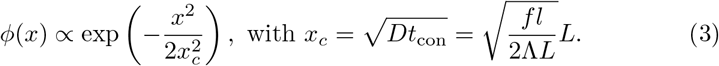

Eq. 3 holds in both one and two spatial dimensions (although recall that in two dimensions we are ignoring corrections that depend on the exact shape of the range) and corresponds to a root mean square distance to the center of 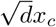. If *x_c_* ≪ *L* then the reproductive value of an individual at the center of the range can be orders of magnitude higher than than one at a typical distance ~ *L*/2 from the center (Fig. 2).

**Figure 2:**
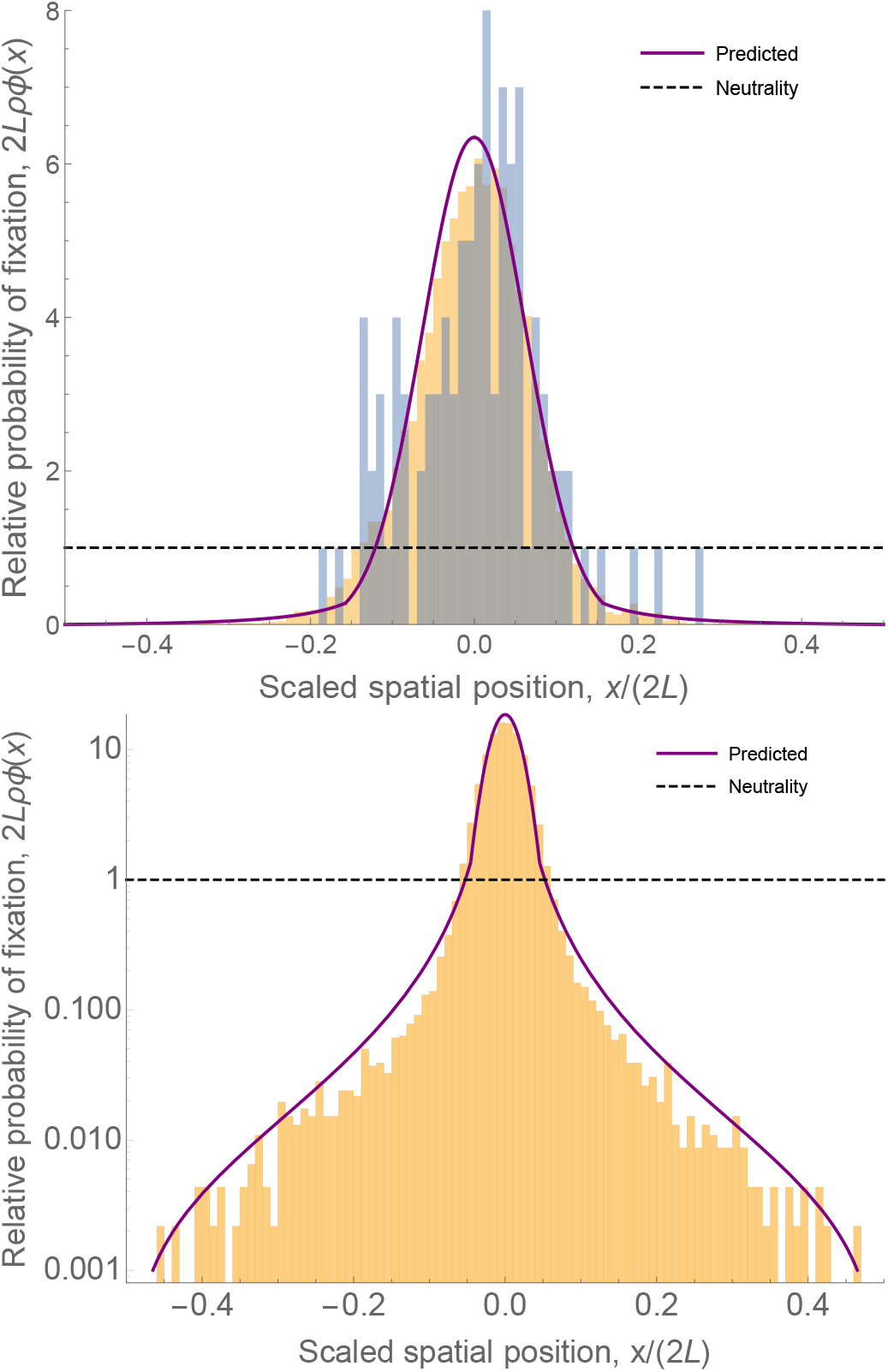
Hitchhiking due to unlinked sweeps concentrates ancestry in the center of the range. The plot shows the probability of that the distant ancestor of a neutral allele was at location *x*, or, equivalently, the fixation probability of a new mutation occurring at *x*. Probabilities are shown on a linear (top) and log (bottom) scale. Histograms show the results of exact forward-time simulations (blue, top panel only) and approximate backward-time simulations (gold). The purple curve shows the predicted distribution: a normal distribution (Eq. 3) inside the center, crossing over to a power law outside (with an additional downturn near the boundaries, Eq. 4), with the crossover between the regimes at the value of *x* at which Eq. 3 and Eq. 4 match. The dashed black line shows a uniform distribution. Parameters for the top panel are *L* = 500, *ρ* = 100, *s* = 0.05, *D* = 0.125, Λ = 0.1, *f* = 1, *K* = 100. Parameters for the bottom panel are as in the right panel of Fig. 1.

From Eq. 3, we see that the ancestral range will be substantially reduced by selection if the rate of sweeps per sexual generation is greater than the ratio of the cline width to the range size: Λ/*f* > *l/L*. It is unclear what ranges these ratios take in natural populations. Λ/(*fK*) is unlikely to be much more than 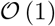 (1) (Weissman and Barton, 2012), but in organisms with many chromosomes (large *K*), Λ/*f* may be substantial. Looking at the right-hand side of the inequality, modeling sweeping alleles by waves spreading across the range necessarily requires *l/L* ≪ 1, so even small values of Λ/*f* may be enough to distort the distribution of ancestry. Surprisingly little is known about typical values of l for the waves of advance of sweeping alleles in nature, but it seems plausible that for many species it should be much smaller than the total species range (Fisher, 1937). For the spread of insecticide resistance in *Culex pipiens* in southern France, the width of the wave of advance was ~ 20 km (Lenormand et al., 1999), much smaller than the global scale of the species range, but the dynamics were more complex than a simple selective sweep (Labbé et al., 2007). Much more is known about the width of stable clines and hybrid zones, which are frequently much smaller than species ranges (Barton and Hewitt, 1985). To the extent that the selection maintaining them is comparable in strength to the selection driving sweeps, these should have roughly the same width as the wavefronts.

### Balance with tightly-linked sweeps

Finding the balance between concentrating effect of unlinked sweeps and the randomizing effect of tightly-linked sweeps is slightly trickier, and we do not know of an exact expression for *ϕ*(*x*). However, we can find an approximate expression by using the fact that the mean squared displacement of the ancestral lineage due to linked sweeps is dominated by rare very tightly-linked sweeps rather than the many loosely-linked ones (Barton et al., 2013). This suggests that for large *x*, the probability that an individual’s ancestor was farther than *x* from the center at time *t*_0_ in the distant past is roughly just the probability that a single very tightly-linked sweep pulled it there at some time within ~ *t*_con_ generations of *t*_0_. Since the distance that a sweep at recombination fraction r pulls the lineage goes like 1/*r*, the rate of sweeps close enough on the genome to pull the ancestry a distance of at least *x* falls off like 1/*x*. Therefore, the probability of finding the ancestry at a distance of at least *x* should also fall off like 1/*x*; the probability density of being exactly at *x*, *ϕ*(*x*)*ρ*(*x*), should then fall off like 1/*x*^2^.

In the Methods, we calculate this more formally, and find:

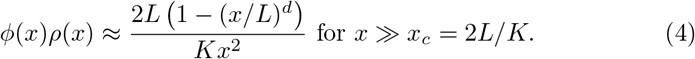

The factor 1 – (*x/L*)^d^ (where *d* = 1 or 2 is the dimension of the habitat) reflects the fact that for very large *x, x ~ L*, most sweeps start at distances less than *x* and cannot pull the lineage that far from the center. For *x* ≪ *x_c_* = 2*L/K*, lineages will tend to experience many sweeps pulling them distances greater than *x* in time ~ *t*_con_, so the approximation used to derive Eq. 4 breaks down; for these small values of *x*, the randomizing effects of moderately-linked sweeps smooth out *ϕ*(*x*) and make it roughly constant.

Barton et al. (2013) describe the randomizing effect of tightly-linked sweeps by “*D*_eff_,” an effective dispersal rate, with 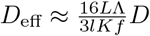 (their Eq. (9)). Comparing Eqs. 3 and 4, however, we see that their effect cannot simply be described as an increase in the dispersal rate, since they create a much longer tail in the spatial distribution of ancestry. Because of this, it is possible that while the bulk of the distribution of ancestry is determined by a balance between unlinked sweeps and dispersal, with linked sweeps too rare to make a difference, linked sweeps make the dominant contribution to the tails of the ancestry distribution (Fig. 2, bottom).

### Combining dispersal and tightly-linked sweeps

Combining Eqs. 3 and 4, we see that unlinked sweeps reduce the effective size of the ancestral range by a factor *x_c_/L*:

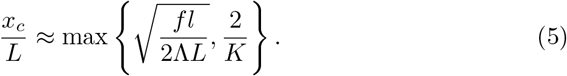

For typical numbers of chromosomes *K*, it would seem that ancestry could be concentrated by about an order of magnitude. However, the result 2/*K* was derived under the assumption that sweeps are distributed uniformly across the genome. If, on the other hand, adaptation is mostly occurring in just a few genes, the rest of the genome will not experience any tightly-linked sweeps, and ordinary dispersal will be the only force counteracting the concentration, meaning that the effect could potentially be much stronger. This has the surprising implication that selection can have a stronger effect on some features of the spatial distribution of ancestry at far-away loci than at those nearby.

### Effect on diversity

While the effect of recurrent sweeps on neutral diversity can be quite large, detecting the effect in data from real populations may be tricky. It might seem to be indistinguishable from a range expansion in the absence of time-series data, but there is a simple way to tell them apart: under recurrent sweeps, there is no serial founder effect reducing diversity away from the center. One way to see this is by looking at isolation by distance. The probability *ψ(x)* that two individuals separated by a distance *x* are genetically identical can be written in terms of the neutral mutation rate *μ* and their coalescence time *T* as

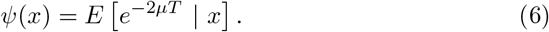

For *x* large compared to the size of a single deme (i.e., the spatial scale over which individuals interact within a generation) and loci far on the genome from any recent sweeps, there are two simple regimes for Eq. 6. If individuals are close together and *μT* ≫ 1, then we expect that the pull due to sweeps is too slow to cause lineages to coalescence before they mutate, and *ψ(x)* is just given by the neutral value, 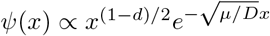 (Barton et al., 2002), which says that the probability of identity falls off rapidly with distance. On the other hand, larger values of *x* are quickly collapsed by the pull of sweeps in time ~ *t*_con_ log(*x/x_c_*), so we expect that *ψ* should be of the form *ψ(x)* ∝ *x*^-2*μt*_con_^. A detailed calculation (see Methods) confirms that this is true for 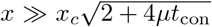; the results are also confirmed by simulations, as shown in Fig. 3. The probability of identity thus has a long tail in distance - if *μt*_con_ is small, individuals at opposite sides of the range (separated by ≈ 2*L*) are nearly as related as individuals separated by, say, *L*/2. Notice that *ψ* does not depend on from where in the range we sampled the pair of individuals. This implies that, while reproductive value is concentrated in the center of the range, genetic diversity is more evenly spread, distinguishing this scenario from a range expansion.

**Figure 3:**
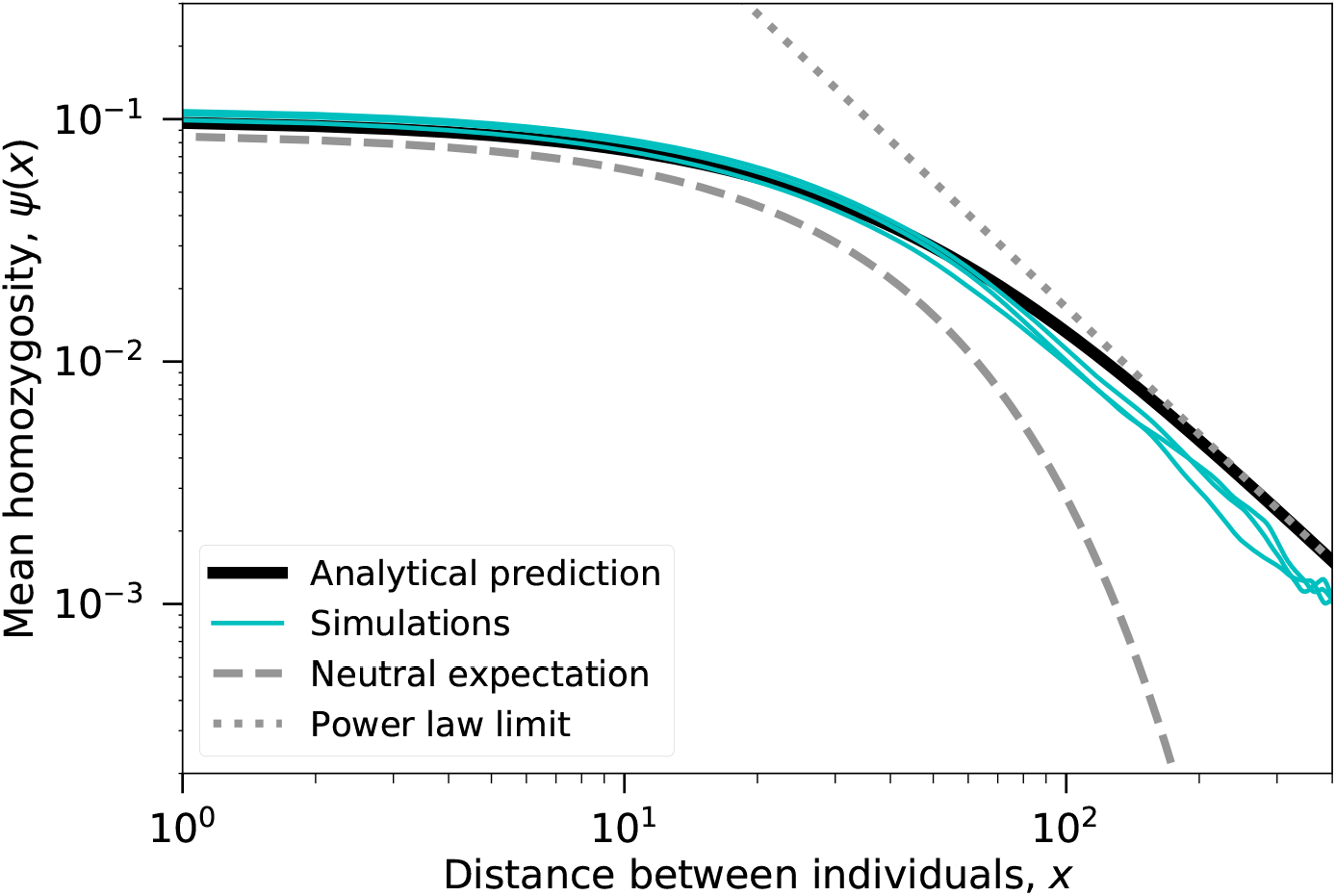
Relatedness between distant individuals has a power-law tail. Expected identity-by-descent *ψ* between a pair of sampled individuals is shown as a function of the distance *x* between them. Cyan curves show the results of three independent forward-time simulations. The solid black curve shows the full analytical prediction, Eq. 25. For large *x*, this approaches a power law, *ψ* ∝ *x*^-2*μt*_con_^ (dotted gray line). This is far higher than it would be in the absence of sweeps, in which case *ψ* would fall off exponentially at rate 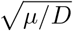 (dashed gray curve). Parameters are as in the left panel of Fig. 1, with *μ* = 3 × 10^-4^ ≈ 1/*t*_con_.

When *x* ≫ *x_c_*, we can approximately invert Eq. 6 to find the distribution of the coalescence time T for one-dimensional ranges. In the Methods, we find that the lineages deterministically approach to within ~ *x_c_* of each other in time ~ *t*_con_ log(*x/x_c_*), after which they coalesce at roughly the same rate as they would in a neutral, well-mixed population of size 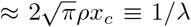. For comparison, in a purely neutral one-dimensional population with strong spatial structure (*L* ≫ *Dρ*), the long-term rate of coalescence is λ_neut_ = *π*^2^*D*/(8*L*^2^) (Maruyama, 1971), so hitchhiking greatly increases the rate of coalescence: λ/λ_neut_ ~ (*L/Dρ*)(*L/x_c_*) ≫ 1. We can also compare the rate of coalescence to that in a well-mixed population with the same pattern of adaptive substitutions. While Barton et al. (2013) showed that spatial structure reduces the coalescence caused by tightly-linked sweeps, for loosely-linked sweeps it can have the opposite effect. In well-mixed populations, unlinked sweeps only substantially increase the rate of coalescence when they become so frequent that they begin to interfere with each other (Λ ~ *f*^2^/*s*) (Weissman and Barton, 2012); for large ranges, coalescence will be increased (i.e., *x_c_* ≪ *L*) at much lower values of Λ than this.

So far in our discussion of diversity, we have ignored loci that are close to recent sweeps. If we are considering large enough loci so that *μt*_con_ ≫ 1, then usually only these recently swept regions will be identical between individuals from different parts of the range. In this case, because each sweep causes coalescence between individuals separated by a large distance *x* over a region of genome with length *r* ∝ 1/*x* (Barton et al., 2013), *ψ* should still have a long tail, but with an exponent that is independent of the population parameters, *ψ* ∝ 1/*x* (see Methods). This characteristic exponent is another effect of rare, tightly-linked sweeps that cannot be accounted for by any effective dispersal rate *D*_eff_.

## Discussion

Because selection and demography are often difficult or impossible to measure directly in natural populations, both are typically inferred from patterns of genetic diversity. This inference can be difficult, because the two processes can produce similar signals. For instance, both purifying selection and population expansion tend to produce site frequency spectra with a relative excess of rare alleles. In order to tease apart the two factors, demography is often first inferred using data from loci that are thought to be neutral, and then the answer is used to infer the pattern of selection at the remaining loci. However, in order for the demography to be inferred correctly, this method requires that the first set of loci be not just neutral, but also unaffected by selection at linked loci. Typically, this is done by using loci that are far from sites where selection is thought to have been important (e.g., Sattath et al. (2011)). Our results suggest that this may be problematic in spatially-structured populations – even diversity at these loci may be strongly affected by unlinked sweeps. Instead, selection and demography should be inferred simultaneously.

### Geometry, not topology

Our results might seem to show that the genetic diversity in a population depends sensitively on the topology of the range and can therefore change dras-tically as the result of small perturbations to the environment. For example, a circular range (which by symmetry has no concentration of ancestry) can be transformed into a linear one (with very concentrated ancestry) by removing a single point. However, this is a misleading interpretation. In fact, a “circular” range is an annulus with radius large compared to its thickness (Fig. 4a). A small perturbation that slightly reduces the population in one part of the range will only have a correspondingly small effect on the distribution of ancestry (Fig. 4b), and the bias of the ancestry ancestry increases smoothly as the perturbation grows (Fig. 4c), until the annulus is completely pinched off (Fig. 4d). More generally, the common-sense intuition that the pattern of diversity should not depend on the details of the shape of the range is correct. All that matters is that, in at least some parts of the range, sweeps are more likely to come from some directions than others. If we consider the vector field defined by the net flow of sweeps, ancestry/reproductive value will tend to concentrate around critical points with positive divergence. (Technically, the distribution of ancestry will evolve according to a convection-diffusion equation.) For the simple range shapes with uniformly distributed sweeps that we have considered in this paper, this occurs in the center of the range. If instead sweeps tended to originate from one end of the range (e.g., if they tend to be introgressed alleles from a hybrid zone), ancestry would concentrate there instead.

**Figure 4:**
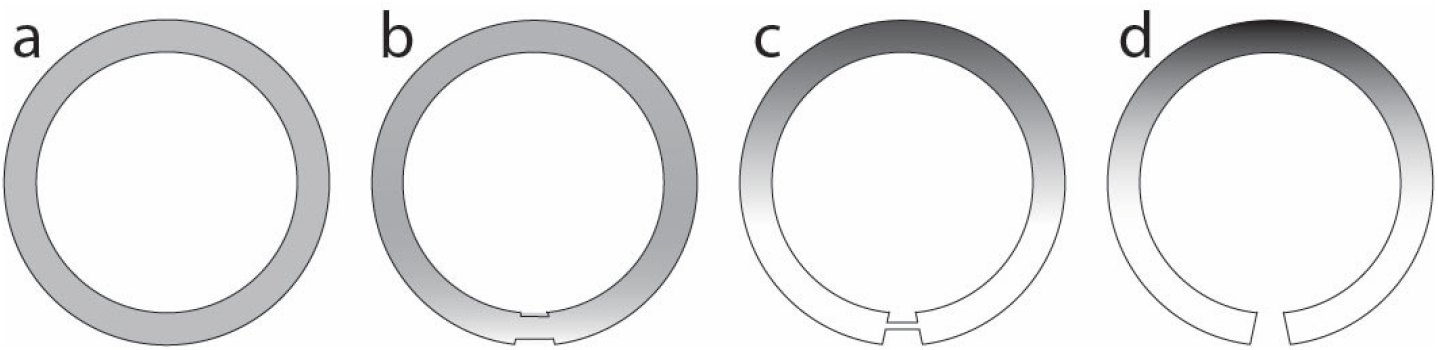
The distribution of ancestry depends smoothly on the shape of the range as it deforms from a perfectly symmetric circle (a) to a curve with endpoints (d). Shading is a schematic representation of the reproductive value of each location, from high (dark) to low (light). In (a), the ancestry is necessarily evenly distributed. Slight asymmetries in the range (b) introduce slight differences in the distribution of ancestry. When the range has a well-defined middle, the ancestry is concentrated there, regardless of whether there is a weak connection between the ends of the range (c) or a strict break (d).

### Extensions

We have focused on a very simple population model. Here we consider several possible modifications. First, we have assumed that the density *ρ* is constant in time. If density fluctuations typically occur on timescales longer than *t*_con_, this approximation should be accurate, and if they are rapid compared to the sweep time *L/c* they should average out, but it is unclear how fluctuations on moderate timescales should interact with dynamics discussed here.

We have also neglected the possibility of rare long-range dispersal. Tightly-linked sweeps already effectively produce occasional long-range jumps in the ancestry of neutral sites, so adding long-range dispersal might not have a large direct effect, but it can have dramatic effects on how sweeps spread (Hallatschek and Fisher, 2014), and therefore a large indirect effect on the hitchhiking dynamics. It is not clear what this effect should be – on the one hand, the sweeps will spread faster, increasing their pull, but on the other hand, the direction of that pull may be less reliably towards the center.

We have also neglected the possibility that many sweeps may be “soft”, starting from multiple alleles (Hermisson and Pennings, 2005), which are likely to be particularly common in spatially-extended populations (Ralph and Coop, 2010). If these alleles typically descend from a recent single ancestor, i.e, are concentrated in a small region at the time when they begin to sweep, then the results should be essentially unchanged, with the possible exception of the coalescent effects of tightly-linked sweeps. The same should be true if sweeps are “firm”, i.e., multiple mutant lineages contribute to each sweep, but the most successful one typically colonizes most of the population. But sweeps in which many widely-spread mutations contribute equally would likely not consistently concentrate ancestry in space.

We have focused on the effect of sweeps on neutral variation, but they will of course also affect selected alleles. Most obviously, if recombination is limited they will interfere with each other (Martens and Hallatschek, 2011). They will interfere even more strongly with weakly-selected variants. We will address these issues in a subsequent manuscript.

## Methods

### Simulations

Forward-time simulations (blue histogram in Fig. 2, left panel in Fig. 1, and Fig. 3) were conducted using the algorithm from Weissman and Barton (2012) (which draws on that of Kim and Stephan (2003)), modified so that population was subdivided into a line of *L* demes of *ρ* individuals each, with random dispersal between adjacent demes. For Figs. 1 and 3, loci were taken to be unlinked (i.e., at a recombination fraction *f*/2 with each other). Because these simulations were extremely computationally demanding, we also conducted approximate backward-time simulations to get better statistics and investigate rare events (right panel of Fig. 1 and gold histograms in 2). These simulations followed a lineage back in time at one neutral locus as it diffused through a continuous one-dimensional space. Sweeps were treated as instantaneous events arising uniformly at random in space and time, with no interference among them. Sweeps occurring at a recombination fraction *r* from the focal locus pulled each lineage an exponentially-distributed distance with mean *c/r* or *c*/(2*r*) (for *r* < *s* and *r* > *s*, respectively), truncated at the origin of the sweep. For the backward time simulations in Fig. 1 and all simulations in Fig. 2, the focal locus was at the center of a linear genome with map length *K* Morgans with sweeps arising uniformly at random across the genome.

### Calculating the “pull” of an loosely-linked sweep

We would like to find the expected spatial displacement of a lineage caused by an loosely-linked sweep, tracing backward in time. To do so, suppose that we sample an allele in a present-day individual in the middle of a very large one-dimensional range, and that a long time ago a selective sweep occurred at a locus a recombination fraction *r* away from the focal allele, starting a very long distance away from our sample. We wish to find the expected location of the ancestor of the sampled allele before the sweep began. Let *p*(*x, τ*) be the probability density for finding the ancestor at location *x τ* generations in the past, with *x* = 0 corresponding to the present location. We want to find:

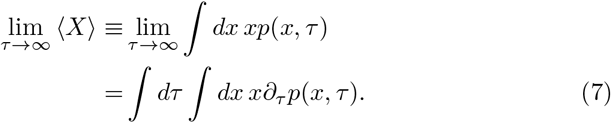

To find *∂_τ_p*, first define *p_i_*(*x, τ*) as the probability density that the ancestor was at location *x* and in genetic background *i*, where *i* = 0 is the ancestral genetic background, and *i* = 1 is the background with the allele that swept. (Note *p* = *p*_0_ + *p*_1_.) If we define *u*(*x, τ*) ≡ *u*_1_(*x, τ*) and *u*_0_(*x, τ*) to be the frequencies of the sweeping allele and the background allele, respectively, with *u*_1_ + *u*_0_ = 1, *p_i_* satisfies the partial differential equation

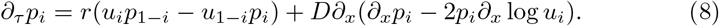

The first term on the right-hand side is the backward-time version of the decay in linkage disequilibrium due to recombination. The second term is backward diffusion; see Appendix A of Hallatschek and Nelson (2008). (Note that their Eq. (3) differs from our Eq. 8 because it includes an additional deterministic drift term due to their use of the co-moving frame of the sweep.) The piece containing *∂_x_* log *u_i_* accounts for the fact that the diffusion is biased towards the direction of increasing frequency of the focal genotype, because migrants of a given genotype are more likely to come from a location where that genotype is frequent than one where it is rare. Technically, in models with discrete generations, Eq. 8 only applies when the recombination rate per generation is small, but we will use it for unlinked loci anyway.

The equivalent of linkage disequilibrium in this system is Δ ≡ *u*_0_*p*_1_ – *u*_1_*p*_0_; we expect it to be small for large *r*. Using Δ to change variables back to *p*, Eq. 8 becomes

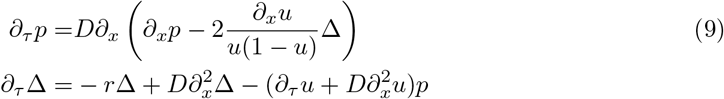

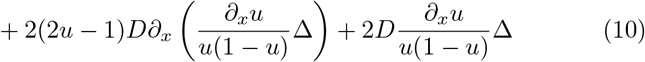

Plugging Eq. 9 into Eq. 7, we have

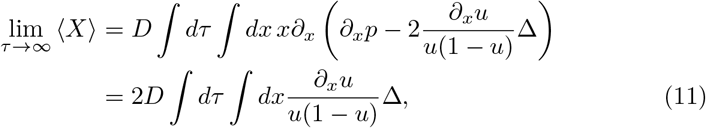

where we have used integration by parts and the fact that *p*(±∞, *τ*) = 0. It now remains to find an expression for Δ. Eq. 10 is quite complicated, but for large r we will have Δ ≪ *p* and the dominant balance will be between the first and third terms on the right-hand side, giving

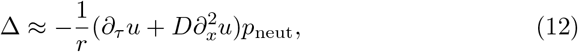

where *p*_neut_ is the value of *p* ignoring the perturbation caused by the sweep, i.e., 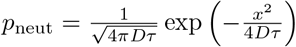. We can simplify this further by noting that u solves Fisher’s equation:

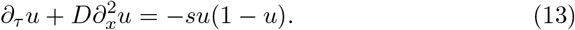

(Recall that *τ* is backward time.) Using this relation and substituting Eq. 12 into Eq. 11, we have

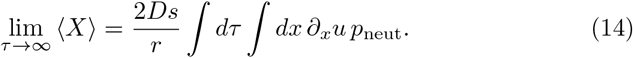

We are interested in the effect of a long-past sweep. Let *τ*_0_ be the time at which the wave of advance passed the point where we sampled the allele; we will take *τ*_0_ to be extremely large. At time *τ*_0_, *p*_neut_ has width 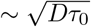, so the wave crosses the region where the ancestor might have lived in a time 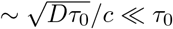, and the integral in Eq. 14 is dominated by times *τ* in the approximate range 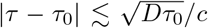. Since *τ* does not vary by much (proportionately) in this interval, *p*_neut_(*x, τ*) ≈ *p*_neut_(*x, τ*_0_) is approximately constant in *τ*. Using this approximation in Eq. 14 yields

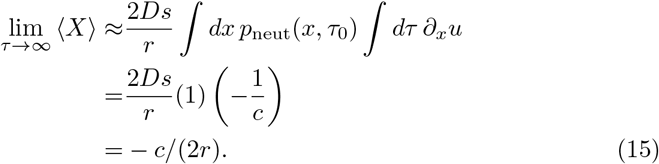

Note that this result did not depend on the form of *p*_neut_, only that it was approximately constant in time; in particular, it also holds if the ancestry settles down to a stationary distribution, as in Eq. 3.

### Effects of noise on sweeps

In Eq. 13 above, we have assumed that sweeps spread as smooth, deterministic waves. In fact, for finite *ρ*, they will be stochastic, and this will tend to reduce their speed *c* (see, e.g., Brunet et al. (2006); Hallatschek and Korolev (2009); and the references in Barton et al. (2013)). We have not attempted a full stochastic derivation of Eq. 15; instead, we simply use the noise-adjusted speed for *c*. In one dimension, this is (Barton et al. (2013), Eq. 5):

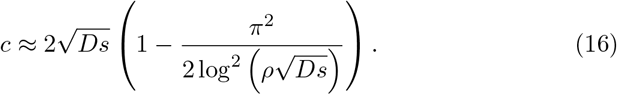

The speed *c* approaches 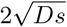 as 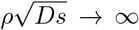, but only very slowly, so the finite-density correction usually cannot be neglected. It is not obvious that substituting Eq. 16 into the final expression Eq. 15 gives the correct answer. We could alternatively, for instance, substitute into the previous line, but this would give the implausible result that the reduction in *c* causes an *increase* in the pull of sweeps. The close agreement between the analytical predictions and simulations in the left panel of Fig. 1 and in Fig. 3 (in which the finite-density correction reduces *c* by approximately 40%) is the best argument that the approach suggested is correct.

### Other kinds of loosely-linked sweep

Above, we have assumed that the sweeping allele spread according to Fisher’s equation, Eq. 13, which describes an allele with a constant selective advantage s. However, the allele may have a varying selective advantage if, for instance, dominance or frequency-dependent effects are important, or if there is environmental variation. More generally, the changing allele frequency is described by

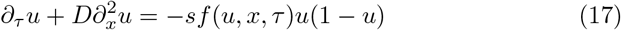

for some function *f*.

Otherwise, the derivation of the expected displacement is the same as above, and we have

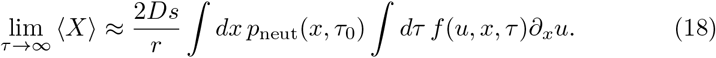

Assuming that *f* is such that *u*(*x,τ*) is still a traveling wave moving at some speed *c*, we can change variables in the second integral to obtain:

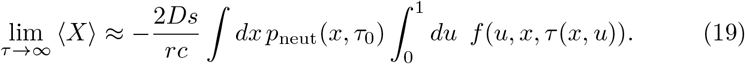

### Effect of tightly-linked sweeps

We wish to calculate *ϕ*(*x*) for large *x*, including the effect of occasional tightly-linked sweeps. It is easiest to consider 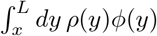, which we can think of as the probability that at some time *t*_0_ in the distant past, the ancestor of a present-day individual was at a distance greater than *x* from the center. For large *x*, we expect that this is dominated by the probability that it was pulled there by a ‘recent’ tightly-linked sweep *t* generations ‘before’ *t*_0_ (i.e., *t* generations closer to the present), with *t* not too large. This sweep must have pulled the lineage out to a distance of at least *xe*^*t/t*_con_^ for it still to be at a distance of at least *x t* generations ‘later’, and therefore the sweep must have originated a distance *z* > *xe*^*t/t*_con_^ from the center. Given that it did, the probability that it pulled the lineage out far enough is exp 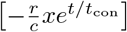. Putting this all together, and using that the density of sweeps per generation per unit map length per distance (or area in two dimensions) at distance *z* from the center and genetic map distance *r* from the focal locus is 2Λ/(*fKL*) (or 4Λ*z*/(*fKL*^2^) in two dimensions), the expected number of sweeps that would have left the lineage more than *x* from the center at time *t*_0_ is:

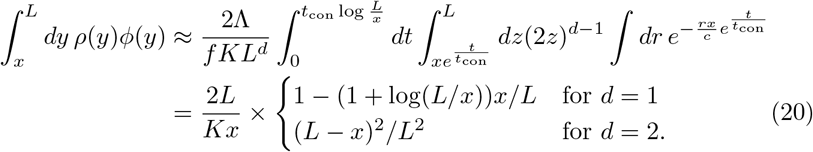

Taking the derivative of both sides of Eq. 20 with respect to *x* gives the probability density, Eq. 4.

Note that Eq. 20 approximates the probability that there is at least one tightly-linked sweep by the expected number of such sweeps, so it is only valid when the right-hand side is small, *x* ≫ 2*L/K*. It also obviously typically breaks down as *x* approaches *L* and the particular geometry of the habitat begins to matter.

### Isolation by distance

We wish to find the probability *ψ(x)* that a pair of lineages a distance *x* apart will be identical at a neutral locus. Let us assume that the locus is far from any recent sweeps. (We relax this assumption below.) Then tracing the ancestry back in time, the separation *X_τ_* between them can be approximated by a Brownian motion, with diffusion constant 2D (since it combines the motion of both lineages), and with the lineages moving together at a mean velocity of ≈ –Λ*cX*/*fL* = –*X/t*_con_ from (unlinked) sweeps that start in between them. In other words, we can approximate the motion by

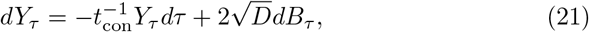

where *B* is a Brownian motion. We write *Y* to emphasize that this is not quite the same as the real path of the lineages *X*. In particular, unlike *X, Y* does not include coalescence. (In two dimensions, *Y* fails to approximate *X* even when the lineages are just very close together, but since most of the coalescence time will be spent at some distance away, it is still a useful approximation.) In addition, Eq. 21 ignores the fact that *X* cannot exceed the diameter of the range 2*L*, and so will only be valid for ranges sufficiently large that lineages are unlikely to bump into the boundaries.

We would like to find an explicit form for Eq. 6. To do this, we can rewrite in terms of the behavior of *Y*. First, note that the rate of coalescence for the two lineages when they are in the same place is 1/*ρ*, and therefore the probability density of coalescence at time *τ* is 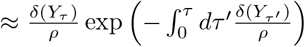, where *δ* is the Dirac delta. (The exponential factor accounts for the possibility that the two lineages have already coalesced.) Plugging this into Eq. 6 gives:

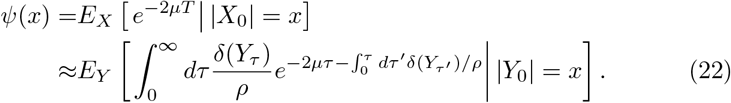

We can use the Feynman-Kac formula (Pham (2009), p25) to rewrite Eq. 22 as an ordinary differential equation:

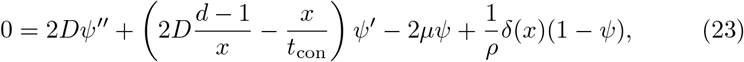

where *δ* is the Dirac delta. Eq. 23 breaks down for *x* → 0 in *d* = 2 dimensions; in this case, some kind of small-scale cutoff is needed, but this does not change the shape of *ψ(x)* at larger scales. In one dimension, to handle the *x* = 0 boundary, we need to understand what we mean by ψ″ and ψ′ at *x* = 0. The correct interpretation is that *x* is actually the *signed* distance between the lineages, i.e., we should remove the absolute value signs around *X*_0_ and *Y*_0_ in Eq. 22 (Barton et al., 2002). Thus 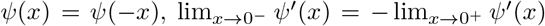, and *ψ*′ has a discontinuity at *x* = 0, i.e., *ψ*″ has a singularity that must cancel with the last term in Eq. 23. This coalescent term can therefore be seen as just a boundary condition that sets the overall normalization of *ψ*. Explicitly, we have:

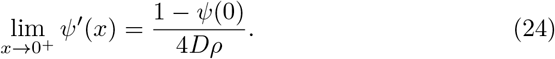

The solution to Eq. 23 can be written exactly in terms of special functions. For *d* = 1 and *x* > 0, Eq. 23 is the Hermite equation, with solution:

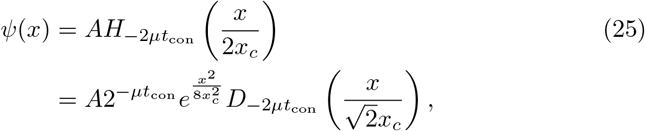

where *H_ν_(z)* is a Hermite function and *D_ν_*(*z*) is a parabolic cylinder function (Wolfram Research (2017) functions `HermiteH` and `ParabolicCylinderD`, respectively), and 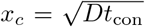. A is a normalization constant, fixed by Eq. 24 to be:

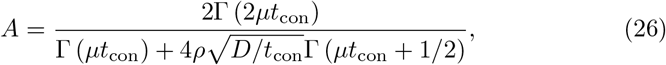

where Γ is the gamma function.

We have not been able to find an exact closed-form expression for the inverse Laplace transform of Eq. 25 (i.e., the distribution of coalescence times) but the mean pairwise coalescent time *τ*_2_ is:

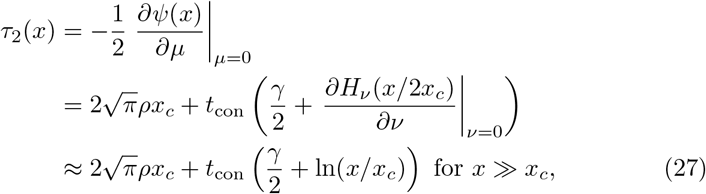

where *γ* ≈ 0.577 is the Euler-Mascheroni constant. Note that two randomly-sampled individuals will typically be a distance ~ *L* apart, so the mean pairwise coalescence time over the whole population can be roughly approximated by *τ*_2_ (*L*).

For large separations 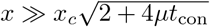, Eq. 25 is approximately:

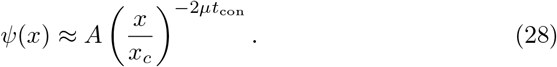

Notice that *ψ* decays only as a power of distance. Up to the normalization constant, Eq. 28 is also valid in two dimensions. For *μt*_con_ ≪ 1, Eq. 26 approaches 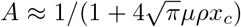, and Eq. 28 approaches the Laplace transform of a simple convolution: first, a nearly deterministic concentration phase lasting *t*_con_ log(*x/x_c_*) generations, followed by an exponentially-distributed phase with mean 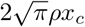, consistent with Eq. 27. In other words, first the lineages are pulled to within ~ *x_c_* of each other, and then undergo neutral coalescence within an effective range of radius ~ *x_c_*.

For *μt*_con_ ≫ 1 and 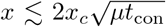, the pull of sweeps is too slow to affect relatedness (by the time the lineages have been pulled together an appreciable distance they will have already mutated), and the solutions to Eq. 23 are close to the neutral solutions in Barton et al. (2002), 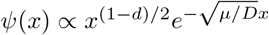 (their Eqs. (10) and (14)).

### Tightly-linked sweeps

Above, we have focused on regions of the genome far from any recent sweeps. Ideally, however, we would like to be able to extend our analysis to include recently-swept regions. As a first approximation, we can say that the main effect of tightly-linked sweeps is that they can cause two widely-separated lineages to rapidly coalesce. The probability that a sweep recombining at rate *r* with the focal neutral locus will cause coalescence between two lineages separated by *x* is ≈ exp(–*rx/c*)/(1 + 2*rϒ*), where *ϒ* is mean coalescence time for two lineages inside the wavefront of the sweep (Barton et al., 2013). We can therefore account for the effect of sweeps uniformly distributed over the genome by changing the coalescence kernel in Eq. 22 from *δ(x)/ρ* to

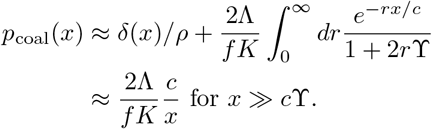

For *x ≫ cϒ*, Eq. 23 then becomes

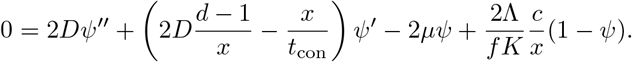

For large *x*, there are two possible tail behaviors for the solution. If 2*μt*_con_ < 1, then the pull of unlinked sweeps is strong enough that it is likely to bring lineages close together before they mutate, and *ψ* ∝ *x*^−2*μt*_con_^ as above. For 2*μt*_con_ > 1, only recently-swept loci share recent enough ancestry to be likely to be identical in distant individuals, and *ψ* ∝ *x*^−1^.

## Acknowledgments

This work would not have been possible without Nick Barton’s generous assistance at all stages.

